# Isolate relatedness is more strongly associated with host and geography compared with phenotypic variation in a broad host range pathogen

**DOI:** 10.64898/2026.11.16.767110

**Authors:** Amit Sharma, Carol L. Groves, Samuel O’Donnell, Richard W. Webster, Horacio D. Lopez-Nicora, Jaime F. Willbur, Damon L. Smith, Emile Gluck-Thaler

## Abstract

Determining the drivers of diversity in plant pathogen populations is a critical for developing effective management strategies. *Sclerotinia sclerotiorum*, a cosmopolitan fungus infecting over 400 plant species, causes considerable yield and economic losses worldwide. However, longstanding questions remain about the association between genetic and phenotypic variation in this pathogen, leaving the drivers of its diversification unresolved. Here, we characterized the diversity of *S. sclerotiorum* isolates from the United States in a global context and tested the hypothesis that phylogenetic relatedness structures phenotypic variation among isolates. Genome-wide variant analysis resolved US isolates into several distinct lineages, with short within-lineage branch lengths consistent with recent common ancestry or clonal propagation. Population structure analysis confirmed two major global clusters, one composed predominantly of Australian isolates and the other largely of isolates from the rest of the world, including the US. Notably, several US isolates grouped within the Australian cluster, suggesting historical intercontinental movement of this pathogen. Contrary to our expectations, isolate relatedness was more strongly associated with data such as host, geographic origin, and year of collection compared with morphological variation, suggesting that phenotypic variation segregates independently of phylogenetic structure. Instead, the distribution of phenotypic variation primarily reflected two trade-offs: rapid radial growth with shorter sclerotial development versus slower growth with prolonged sclerotial development, and the production of many small sclerotia versus fewer larger ones. Together, our results shed light on the complex relationships between phenotypic, ecological, and phylogenetic variation and bring us one step closer to predicting how diversity is structured in natural populations of a global fungal pathogen.

## Introduction

*Sclerotinia sclerotiorum* is a formidable and economically significant fungal plant pathogen because of its extraordinarily broad host range and complex lifestyle (McCaghey et al. 2019). This cosmopolitan pathogen has been documented in nearly all major agricultural regions and is capable of causing highly destructive white mold or stem rot diseases in a wide variety of economically important crops, including soybean (*Glycine max*), sunflower (*Helianthus annus),* canola (*Brassica napus*), lettuce (*Lactuca sativa*), carrot (*Daucus carota*), and potato (*Solanum tuberosum*) (Boland and Hall 1994; Willbur et al. 2019). Management of diseases caused by *S. sclerotiorum* is challenging because complete host resistance has not been identified in most major susceptible crops and only partial, quantitatively inherited resistance governed by multiple genes is available (Bolton et al. 2006; McCaghey et al. 2019). The development of durably resistant cultivars is contingent upon resistance evaluations that adequately capture the full breadth of genotypic and phenotypic diversity present in *S. sclerotiorum* populations. Resistance identified and validated against a limited set of *S. sclerotiorum* isolates risks rapid breakdown upon deployment in the field, where the crop is exposed to far more genetically diverse pathogen populations than those used in controlled resistance screening (Willbur et al. 2017). It is therefore imperative that the diversity of *S. sclerotiorum* populations be thoroughly characterized, and the origins of this variation elucidated.

The reproductive biology of *S. sclerotiorum* shapes its genetic structure at a population level. Central to the reproductive biology of this pathogen is the asexual production of sclerotia, which are compact, melanized masses of aggregated mycelium that function as recalcitrant survival structures capable of remaining viable in the soil for up to eight years (Willbur et al. 2019). Under favorable environmental conditions, sclerotia germinate through one of two distinct pathways: myceliogenic germination, in which sclerotia directly produce vegetative mycelium that subsequently forms new sclerotia, perpetuating the asexual cycle; or carpogenic germination that initiates the sexual cycle, in which sclerotia develop apothecia that release ascospores, which serve as the primary inoculum for initiating new infections (Bolton et al. 2006; Sharma et al. 2025; Willbur et al. 2019). As a homothallic fungus, *S. sclerotiorum* harbors both mating types within the same individual, conferring the capacity for self-fertilization and reducing the likelihood of outcrossing. This limited potential for outcrossing is further compounded by the monocyclic nature of the pathogen, which completes only one life cycle per growing season, limiting opportunities for genetic recombination. Moreover, even when two isolates initiate the sexual cycle in proximity, successful genetic exchange is dependent on compatibility, and compatible isolates tend to be genetically similar, further diminishing the probability of generating novel genotypes distinct from the parental types (Buchwaldt et al. 2022). Given these constraints on the generation of genetic variation, one might reasonably predict that *S. sclerotiorum* populations would be dominated by clonal lineages with limited genetic recombination and minimal within-population diversity (Buchwaldt et al. 2022). Indeed, many early studies relying on mycelial compatibility groupings (MCGs) and restriction fragment length polymorphism (RFLP) analyses reported low genetic variability and a predominantly clonal population structure across regions including Australia, Canada, and the US (Attanayake et al. 2019; Cubeta et al. 1997; Ekins et al. 2011; Kohli et al. 1995).

However, contrary to what early evidence suggested, recent sequencing approaches have provided compelling evidence of genetic recombination in natural populations, demonstrating that outcrossing does occur frequently in *S. sclerotiorum* (Atallah et al. 2004; Attanayake et al. 2014, 2019; Derbyshire et al. 2019a, 2025). Additional sources of genetic variation include point mutations and transposable element activity (Buchwaldt et al. 2022; Derbyshire et al. 2025). Genetic diversity is geographically structured through multiple dispersal mechanisms, including the movement of sclerotia via contaminated seeds and farm machinery within and between fields (Anderson and Kohn 1995), and the aerial transport of ascospores by wind (Cubeta et al. 1997). Despite isolation by distance effects, inoculum dispersal ensures the continuous redistribution of genetically diverse isolates across agricultural landscapes, posing a significant challenge to durable disease management because greater diversity may facilitate more rapid evolution of virulence and fungicide resistance over time (Derbyshire et al. 2019b). Additionally, genetically diverse populations may differ in their sensitivity to biocontrol agents, further complicating the implementation of integrated disease management strategies (Sharma et al. 2025).

Achieving a comprehensive understanding of the *S. sclerotiorum* population diversity is a critical first step towards the design of effective and durable disease management strategies. However, the structure of genetic and phenotypic diversity in *S. sclerotiorum* populations across the US and their underlying drivers remain poorly understood. The objectives of this study were therefore to: (1) characterize the genetic diversity of *S. sclerotiorum* isolates collected from various regions across the US and place it in a global context to determine how local and regional populations are structured and how they relate to the broader global population; (2) evaluate morphological phenotypic variation among US isolates; and (3) investigate whether patterns of genetic diversity are more strongly associated with phenotypic traits compared with ecological variables including host of origin, geographic location, and year of collection, with the overarching aim of determining the nature of the association between genetic and phenotypic variation in this pathogen.

## Materials and Methods

### Sample collection

A total of 114 isolates of *S. sclerotiorum*, originally collected from across the United States, were selected for short-read sequencing based on host plant, field location, and year of isolation from culture collections maintained at the University of Wisconsin–Madison, Madison, WI; North Dakota State University, Fargo, ND; and The Ohio State University, Columbus, OH, along with one isolate each from Canada, Poland, and Chile (Table 1). DNA sequencing data from an additional 102 representative *S. sclerotiorum* isolates were downloaded from NCBI BioProject PRJNA1120954. The isolates with publicly available data were selected based on their geographical location, including Australia, Canada, Chile, Norway, Poland, and the United Kingdom (Fig. 1A). A *S. minor* isolate SS60, originally collected from Poland and maintained in the Smith lab culture collection at the University of Wisconsin–Madison, was used as an outgroup.

**Figure. 1.**
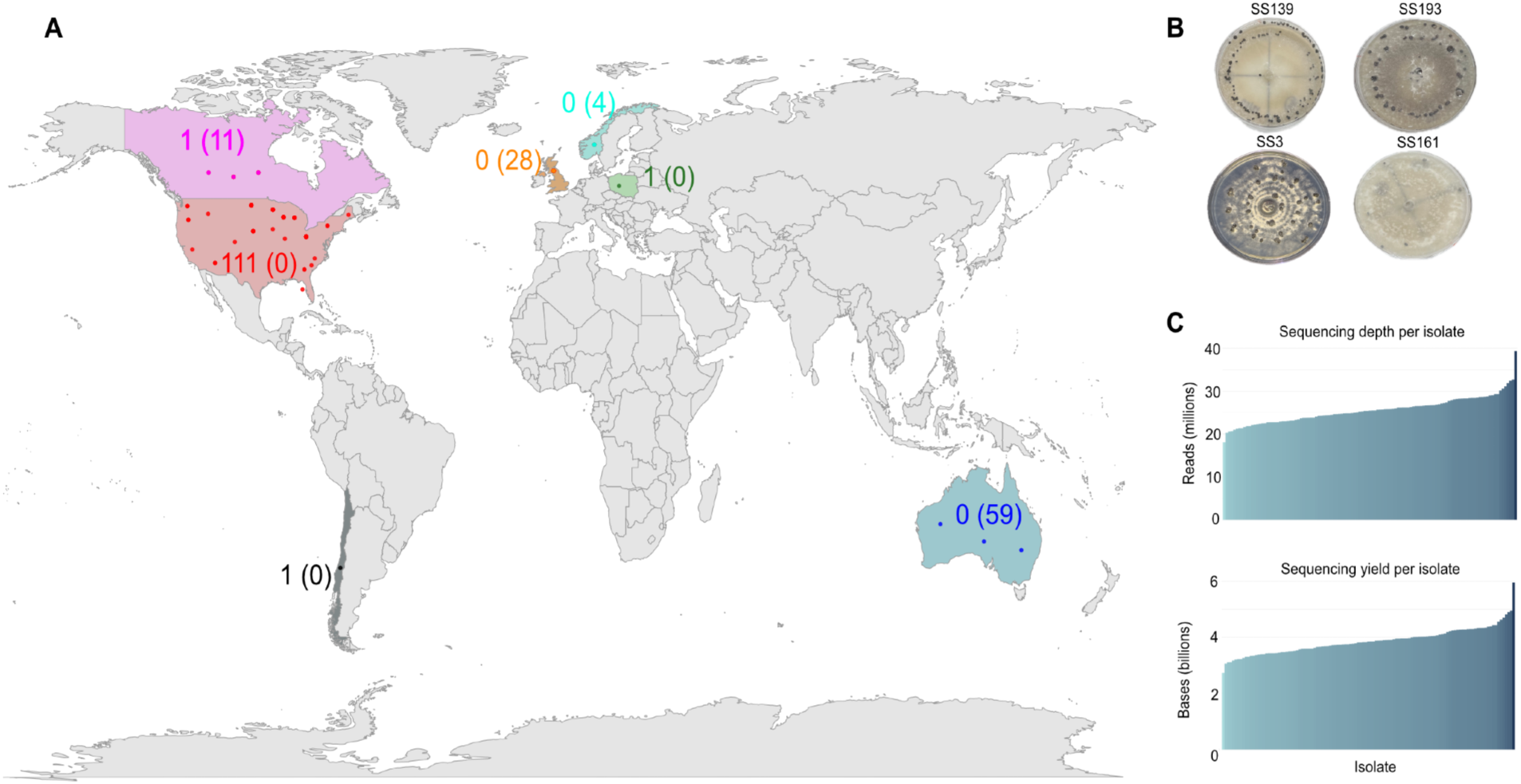
**(A)** Geographic distribution of *Sclerotinia sclerotiorum* isolates used in this study, including newly sequenced (n=114) and those with publicly available data (n=102). For each location, numbers outside parentheses indicate newly sequenced isolates, while numbers in parentheses represent isolates whose data were obtained from NCBI. Isolates were distributed as follows: United States 111 (0), Canada 1 (11), United Kingdom 0 (28), Norway 0 (4), Poland 1 (0), Chile 1 (0), Australia 0 (59). **(B)** Example cultures of *Sclerotinia sclerotiorum* isolates showing colony morphology (SS139, SS193, SS3, and SS161) on potato dextrose agar (PDA) at 21°C for 7 days. Sequencing depth and yield metrics for *Sclerotinia sclerotiorum* isolates.

**Table 1.** *Sclerotinia sclerotiorum* isolates selected for short-read sequencing. A total of 114 isolates were obtained from culture collections maintained at the University of Wisconsin–Madison (Madison, WI), North Dakota State University (Fargo, ND), and The Ohio State University (Columbus, OH), but were originally collected from locations across the United States, and one each from Canada, Poland, and Chile. Host plant, collection location, and year of isolation are listed for each isolate.

| Serial No. | isolate | host | location | year |
|---|---|---|---|---|
| 1 | SS10 | Soybean | Michigan | Unknown |
| 2 | SS111 | Soybean | Chile | 2018 |
| 3 | SS113 | Vinca (periwinkle) | Wisconsin | 2019 |
| 4 | SS114 | Soybean | Wisconsin | 2021 |
| 5 | SS121 | Unknown | Wisconsin | 2021 |
| 6 | SS122 | Unknown | Wisconsin | 2022 |
| 7 | SS123 | Rapeseed | Unknown | 2022 |
| 8 | SS137 | Potato | Michigan | 2019 |
| 9 | SS139 | Potato | Michigan | 2019 |
| 10 | SS141 | Potato | Michigan | 2019 |
| 11 | SS142 | Potato | Michigan | 2019 |
| 12 | SS143 | Potato | Michigan | 2023 |
| 13 | SS144 | Potato | Michigan | 2023 |
| 14 | SS145 | Potato | Michigan | 2023 |
| 15 | SS146 | Potato | Michigan | 2023 |
| 16 | SS149 | Dry bean | Nebraska | 2023 |
| 17 | SS15 | Soybean | Wisconsin | 2000 |
| 18 | SS150 | Soybean | Ohio | 1999 |
| 19 | SS151 | Soybean | Ohio | 2015 |
| 20 | SS152 | Soybean | Ohio | 2015 |
| 21 | SS153 | Soybean | Ohio | 2015 |
| 22 | SS154 | Soybean | Ohio | 2015 |
| 23 | SS155 | Soybean | Ohio | 2015 |
| 24 | SS156 | Soybean | Ohio | 2015 |
| 25 | SS157 | Soybean | Ohio | 2015 |
| 26 | SS158 | Soybean | Ohio | 2015 |
| 27 | SS159 | Soybean | Ohio | 2015 |
| 28 | SS16 | Soybean | Unknown | Unknown |
| 29 | SS160 | Soybean | Ohio | 2015 |
| 30 | SS161 | Soybean | Ohio | 2015 |
| 31 | SS162 | Soybean | Ohio | 2015 |
| 32 | SS163 | Soybean | Ohio | 2015 |
| 33 | SS164 | Soybean | Ohio | 2015 |
| 34 | SS165 | Soybean | Ohio | 2015 |
| 35 | SS166 | Soybean | Ohio | 2015 |
| 36 | SS167 | Soybean | Ohio | 2015 |
| 37 | SS168 | Soybean | Ohio | 2015 |
| 38 | SS169 | Soybean | Ohio | 2015 |
| 39 | SS170 | Soybean | Ohio | 2015 |
| 40 | SS171 | Soybean | Ohio | 2015 |
| 41 | SS172 | Soybean | Ohio | 2021 |
| 42 | SS173 | Soybean | Ohio | 2022 |
| 43 | SS174 | Soybean | Ohio | 1999 |
| 44 | SS175 | Soybean | Ohio | 2023 |
| 45 | SS176 | Soybean | Ohio | 2023 |
| 46 | SS177 | Soybean | Ohio | 2023 |
| 47 | SS178 | Soybean | Ohio | 2015 |
| 48 | SS179 | Soybean | Ohio | 2015 |
| 49 | SS180 | Dry bean | North Dakota | 2008 |
| 50 | SS181 | Dry bean | Canada | 2008 |
| 51 | SS182 | Dry bean | North Dakota | 2008 |
| 52 | SS183 | Canola | North Dakota | 2007 |
| 53 | SS184 | Soybean | North Dakota | 2008 |
| 54 | SS185 | Dry bean | North Dakota | 2008 |
| 55 | SS186 | Dry bean | North Dakota | 2008 |
| 56 | SS187 | Dry bean | North Dakota | 2008 |
| 57 | SS188 | Soybean | North Dakota | 2008 |
| 58 | SS189 | Dry bean | Minnesota | 2008 |
| 59 | SS19 | Soybean | Unknown | 2012 |
| 60 | SS190 | Sunflower | Minnesota | 2008 |
| 61 | SS191 | Soybean | Minnesota | 2008 |
| 62 | SS192 | Dry bean | Nebraska | 2008 |
| 63 | SS193 | Dry bean | North Dakota | 2008 |
| 64 | SS194 | Canola | North Dakota | 2008 |
| 65 | SS195 | Sunflower | North Dakota | 2008 |
| 66 | SS196 | Sunflower | North Dakota | 2008 |
| 67 | SS197 | Canola | North Dakota | 2008 |
| 68 | SS198 | Sunflower | North Dakota | 2008 |
| 69 | SS199 | Soybean | North Dakota | 2008 |
| 70 | SS20 | Soybean | Unknown | 2012 |
| 71 | SS200 | Soybean | North Dakota | 2008 |
| 72 | SS201 | Sunflower | North Dakota | 2008 |
| 73 | SS202 | Canola | Colorado | 1992 |
| 74 | SS203 | Dry bean | Nebraska | 2008 |
| 75 | SS204 | Soybean | Ohio | 2002 |
| 76 | SS205 | Soybean | Iowa | 2008 |
| 77 | SS206 | Soybean | Wisconsin | 2000 |
| 78 | SS207 | Soybean | Wisconsin | 2000 |
| 79 | SS208 | Soybean | Wisconsin | 2002 |
| 80 | SS209 | Canola | North Dakota | 2002 |
| 81 | SS210 | Soybean | North Dakota | 2008 |
| 82 | SS211 | Lentil | Washington | Unknown |
| 83 | SS212 | Pea | Washington | Unknown |
| 84 | SS213 | Gourd | Washington | Unknown |
| 85 | SS214 | Potato | Washington | Unknown |
| 86 | SS215 | Lettuce | California | Unknown |
| 87 | SS216 | Cauliflower | Oregon | Unknown |
| 88 | SS217 | Snap bean | Oregon | Unknown |
| 89 | SS218 | Parsley | South Carolina | Unknown |
| 90 | SS219 | Cabbage | North Carolina | Unknown |
| 91 | SS220 | Dry bean | Idaho | Unknown |
| 92 | SS221 | Canola | Arizona | Unknown |
| 93 | SS222 | Cabbage | Georgia | Unknown |
| 94 | SS223 | Carrot | Georgia | Unknown |
| 95 | SS224 | Potato | Florida | Unknown |
| 96 | SS225 | Lettuce | Arizona | Unknown |
| 97 | SS226 | Bush bean | Maine | Unknown |
| 98 | SS227 | Canola | New York | Unknown |
| 99 | SS228 | Canola | New York | Unknown |
| 100 | SS229 | Dry bean | North Dakota | 2023 |
| 101 | SS23 | Soybean | Wisconsin | 2003 |
| 102 | SS26 | Soybean | Wisconsin | 2012 |
| 103 | SS27 | Soybean | Wisconsin | Unknown |
| 104 | SS28 | Soybean | Wisconsin | Unknown |
| 105 | SS3 | Soybean | Wisconsin | 2012 |
| 106 | SS30 | Soybean | Wisconsin | 2012 |
| 107 | SS43 | Canola | Nebraska | Unknown |
| 108 | SS44 | Greenbean | Wisconsin | 2009 |
| 109 | SS45 | Petunia | Minnesota | 2009 |
| 110 | SS46 | Soybean | Minnesota | 2013 |
| 111 | SS5 | Soybean | Illinois | Unknown |
| 112 | SS62 | Rapeseed | Poland | 2013 |
| 113 | SS71 | Soybean | Wisconsin | 2014 |
| 114 | SS72 | Soybean | Wisconsin | 2014 |

### DNA extraction and sequencing of fungal isolates

DNA was extracted from 114 isolates of *S. sclerotiorum* and one *S. minor* isolate maintained in the Smith lab collection. Sclerotia from each isolate were surface disinfested in 95% ethanol for 60 seconds, rinsed in sterile deionized water, treated with 0.825% sodium hypochlorite for 60 seconds, and again rinsed in sterile deionized water, then were blotted dry on sterile filter paper (Qualitative P8, porosity coarse; Fisher Scientific). The sclerotia were then bisected and placed onto potato dextrose agar (PDA; Difco^TM^) in 100-× 15-mm Petri plates and incubated at 21°C (Sharma et al. 2026a). Examples of cultures of *S. sclerotiorum* are shown in Fig. 1B. A 5-mm mycelial plug of the fungal isolate from the leading edge of the actively growing culture was transferred to a Petri plate containing sterile potato dextrose broth (PDB; Difco™). After 4-5 days of incubation, the resulting mycelial mat was harvested by vacuum filtration and stored at −20°C until further use. Genomic DNA was extracted from mycelia using the FastDNA SPIN Kit and the FastPrep Instrument following the manufacturer’s instructions. The extracted DNA was then diluted to 750 ng in a final volume of 25 µL and used for whole genome sequencing (WGS). Paired end reads of the genomes of all isolates were generated using the Illumina sequencing platform at the University of Wisconsin–Madison Biotechnology Center, Madison, WI. Raw sequencing data were processed to assess data quality, with sequencing depth ranging from 20-40 million reads per isolate and total sequencing yield of 3-6 billion bases per sample (Fig. 1C).

### Variant calling

We used WGS data from 114 newly sequenced isolates of the US collection together with publicly available short-read genomic data from 102 isolates retrieved from NCBI for variant calling. Single Nucleotide Variants (SNVs) were called across all genomes using the snpArcher workflow executed with Snakemake version 9.13.2 (https://github.com/harvardinformatics/snpArcher) (Mirchandani et al. 2024). Raw reads were aligned to the reference genome (*Sclerotinia sclerotiorum* 1980 UF70, NCBI BioProject PRJNA15530) using the Burrows-Wheeler Aligner (BWA) (Li 2013). Following alignment, we used Genome Analysis Toolkit (GATK) HaplotypeCaller, which models local haplotypes to improve SNV detection accuracy to call variants independently for each sample (McKenna et al. 2010). Per-sample variant calls were generated in genomic VCF (gVCF) format and subsequently combined across all samples through joint genotyping to ensure consistent variant representation and genotyping across the dataset. Jointly genotyped variants were subjected to quality filtering to retain high-confidence sites, resulting in a multi-sample VCF containing high-confidence SNVs for downstream analyses (Van der Auwera et al. 2013). Default parameters were used as implemented in the snpArcher pipeline unless otherwise specified.

### Phylogenetic analysis

We focused our phylogenetic analysis on a subset of the jointly genotyped variant dataset corresponding to newly sequenced isolates from the US collection. To generate a multiple sequence alignment, we converted the multi-sample VCF into a concatenated SNV alignment using vcf2aln tool (Parker et al. 2020) (https://github.com/campanam/vcf2aln). The resulting alignment was exported in FASTA format and used for maximum-likelihood (ML) phylogenetic inference in IQ-TREE2 under the general time reversible (GTR) substitution model (Minh et al. 2020). Because the alignment contained only variable sites, ascertainment bias correction (+ASC) was applied during model fitting. Branch support was assessed using the ultrafast bootstrap approximation (UFBoot) and the Shimodaira-Hasegawa approximate likelihood ratio test (SH-aLRT), each with 1000 replicates (Minh et al. 2020). The maximum-likelihood tree was inferred by log-likelihood maximization, and branch support values were mapped onto the final tree. Phylogenetic trees were inferred both with and without an outgroup. For the rooted analysis, *S. minor* isolate SS60 was included as an outgroup to root the tree, while an additional unrooted tree was inferred excluding the outgroup to assess relationships among the ingroup isolates.

### Population differentiation analyses

We filtered the variant data from all *S. sclerotiorum* isolates (newly sequenced and publicly available) and converted them to PLINK binary format using PLINK v1.9, retaining only biallelic SNVs composed of A, C, G, or T alleles (Chang et al. 2015). Variants with >10% missing genotypes were excluded. A maximum chromosome count of 50 was specified to accommodate the fragmented (non-telomere-to-telomere) reference assembly, which consists of multiple scaffolds rather than complete chromosomes. To reduce the confounding effects of linkage disequilibrium, SNPs were pruned using a sliding window approach (window size = 50 SNPs, step size = 10 SNPs, r² threshold = 0.1). The resulting pruned dataset was used for all subsequent population structure analyses. STRUCTURE analyses were performed using fastSTRUCTURE framework (Raj et al. 2014). Models were run for K = 2 to K = 5 ancestral populations, with five independent replicates per K value to assess consistency across runs. Output files containing individual ancestry proportions (Q matrices) were collected for downstream visualization and comparison. Q-matrices were post-processed in R to harmonize cluster labels across replicates, and average pairwise similarity among runs was calculated for each K.

To investigate relationships among *S. sclerotiorum* isolates, we constructed a phylogenetic network using SplitsTree (version 6.6.2) (Huson and Bryant 2024). The filtered multi-sample VCF was converted to a PHYLIP-format SNP alignment using vcf2phylip. The resulting PHYLIP file was imported into SplitsTree for network construction. Within SplitsTree, pairwise genetic distances were calculated using p-distances. Phylogenetic networks were inferred using the NeighborNet algorithm, as implemented in SplitsTree, to visualize conflicting phylogenetic signals and reticulate relationships among samples.

We estimated within population nucleotide diversity (π) as the mean pairwise distance among clone-corrected isolates within each population. Between population divergence was summarized as the mean pairwise distance between populations (d_xy_) and as Nei’s net divergence, which removes the contribution of ancestral within-population diversity. A distance-based F_ST_ was computed as D_a_/d_xy_. A 95% confidence interval for D_a_ was obtained by bootstrap resampling of isolates within each population (1,000 replicates). Genetic variance was partitioned within and among the two populations by distance-based analysis of molecular variance (AMOVA) on the clone-corrected squared-distance matrix, and the fixation index Φ_ST_ was calculated as the ratio of the among population variance component to the total. Divergence statistics were calculated on the clone-corrected matrix with a custom function popstats in R (Skoglund et al. 2015). AMOVA was performed using the amova function in the R package pegas (Paradis 2010).

### Phenotypic characterization

Prior to starting experiments, we mass produced sclerotia of the US isolates by transferring agar plugs from an actively growing culture of each respective isolates to prepared carrot discs in 250-mL Erlenmeyer flasks (Willbur et al. 2017). After 30 days of incubation in the dark at room temperature, sclerotia were collected by passing the carrot culture through a 2.00 mm sieve (US standard, American Scientific) to remove the plant material. The sclerotia were rinsed thoroughly and dried at room temperature for 48 hours prior to being stored at 4°C in sterile 50 mL tubes.

Sclerotia were subsequently retrieved, surface-disinfested, bisected, and placed onto PDA plates at 21°C, as described previously. The phenotypic characteristics of isolates were evaluated by transferring a 5-mm agar plug from the leading edge of actively growing culture onto PDA in 100-× 15-mm Petri plates and incubating them at 21°C under a 12-hour photoperiod. The whole procedure was performed three times with three replicates for each isolate. Radial mycelial growth was measured daily along two perpendicular axes (x and y) of each plate until the whole agar surface was fully covered with mycelia (Sharma et al. 2026a). Standardized area under the growth curve (stAUGC) was calculated and used for statistical analyses. The stAUGC was computed using the same approach as the standardized area under the disease progress curve (Campbell and Madden 1990; Sharma et al. 2026a; Willbur et al. 2017). Data on phenotypic traits including number of sclerotia produced per plate, sclerotial diameter, sclerotial weight, and number of days to initiation of sclerotia formation and sclerotia maturation were recorded for all isolates. Sclerotial initiation was defined by the aggregation of hyphae into compact masses, while maturation was characterized by consolidation and melanin deposition in the internal rind cells (Bolton et al. 2006; Townsend and Willetts 1954). We performed principal component analysis (PCA) on all six above-mentioned phenotypic traits for all *S. sclerotiorum* isolates from the US collection. Prior to analysis, all traits were standardized (centered and scaled) to account for differences in measurement units and ranges. PCA was conducted using the prcomp() function in R (Groth et al. 2013); R Core Team, 2025). A biplot was generated to simultaneously visualize isolate distribution in principal component space and trait loadings as vectors,indicating the direction of each trait contribution to the principal components.

### Effect of host, location, and year on phenotypic traits

To examine relationships among the phenotypic variables, we performed correlation analyses. Pearson’s correlation coefficients (r) and corresponding *P*-values were calculated using the rcorr() function from the Hmisc package in R (Harrell Jr 2003). Correlations were considered statistically significant at *P* < 0.01.

Based on the correlation analysis, a subset of quantitative traits, including median stAUGC and median sclerotial weight, was retained for each isolate to evaluate the effects of host, geographic location, and year of collection. To ensure comparisons and reduce bias associated with sparsely represented categories, datasets were filtered prior to analysis. For each categorical variable (host, location, and year), only groups represented by three or more isolates were retained. Isolates with missing or unknown data were excluded from further analyses. Phylogenetic linear mixed-effects models were fitted using the pglmm function in the phyr package in R (Li et al. 2020); R Core Team, 2025), and analysis of variance was used to assess the effects of host, geographic location, and year on these quantitative phenotypic traits. Separate models were fitted for each explanatory variable, which was treated as a fixed effect, while phylogeny and the remaining categorical variables were included as random effects to account for shared variance among isolates. When the fixed effect in a given model was significant (*P* < 0.05), we calculated estimated marginal means and conducted pairwise comparisons among factor levels. These pairwise comparisons were interpreted using Fisher’s LSD and significant differences among levels were summarized using compact letter displays generated with the multcompView package (Graves et al. 2015).

### Phylogenetic signal analyses

Phylogenetic signal analyses were conducted using Blomberg’s K and Pagel’s λ to assess the association between phylogeny and the measured phenotypic traits for newly sequenced isolates from the US collection (Münkemüller et al. 2012). Each index was tested against the null hypothesis of no phylogenetic signal, in which trait values are randomly distributed across the phylogeny. Blomberg’s K and Pagel’s λ for numerical traits were estimated in R using the phylosig function in the phytools package (Revell 2024). Phylogenetic signal in the categorical traits host, location, and year was assessed by fitting maximum-likelihood Mk models of discrete trait evolution using the fitDiscrete function in the geiger R package, with Pagel’s λ estimated as a tree-transformation parameter via likelihood optimization (Pennell et al. 2014). Blomberg’s K and Pagel’s λ values range from 0 to 1. Values near 0 indicate little to no phylogenetic signal, suggesting that traits vary independently of phylogenetic structure, and that closely related isolates are not more phenotypically similar than distantly related ones. Values >0 and <1 indicate partial phylogenetic dependence, whereas values approaching 1 indicate strong phylogenetic signal, with closely related isolates exhibiting similar trait values (Tuya et al. 2024).

## Results

### Genetic diversity

Two phylogenetic trees were inferred from the SNV data, one including the *S. minor* isolate SS60 as an outgroup to root the phylogeny and assess evolutionary divergence (Supplementary Fig. 1A), and a second without the outgroup to improve resolution of relationships among closely related isolates (Fig. 2A). The maximum likelihood tree showed evolutionary distances ranging up to 0.06 substitutions per site. Isolates clustered into several distinct lineages, with the majority forming tight clusters characterized by short branch lengths (<0.01 substitutions per site), suggesting recent common ancestry or potential clonal relationships. Most terminal and shallow internal nodes were strongly supported, with ultrafast bootstrap (UFBoot) and SH-like aLRT values frequently ≥95. To investigate potential host or geographic population patterns, isolates were color-coded by host plant (Supplementary Fig. 1B) and collection location (Supplementary Fig. 1C).

**Figure. 2.**
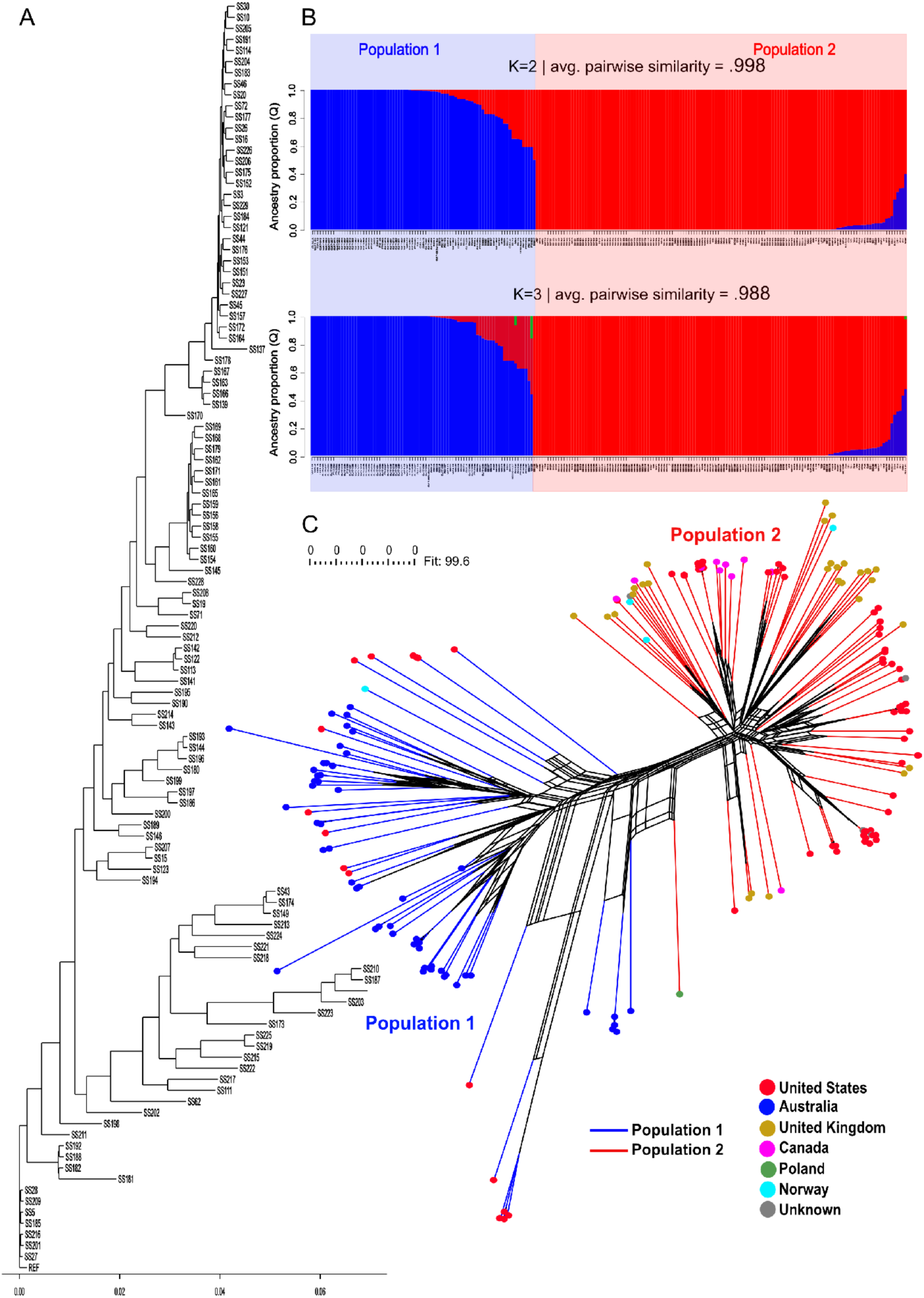
**(A)** Maximum-likelihood phylogenetic tree of newly sequenced *Sclerotinia sclerotiorum* isolates based on whole genome-wide SNP analysis. Phylogeny constructed after removal of SS60 (outgroup) to improve resolution of relationships among closely related isolates. Branch lengths are proportional to genetic distance. **(B)** Population structure inferred from genome-wide SNP data using fastSTRUCTURE at K = 2 and K = 3. Each vertical bar represents an individual isolate, with colors indicating proportional ancestry (Q) assigned to each genetic cluster. At K = 2, two distinct population clusters were observed (blue, Population1; red, Population 2), with average pairwise similarity = 0.998. At K = 3, a third cluster emerges at low ancestry proportions in a small subset of individuals (green, Population 3), accompanied by a slight decrease in clustering consistency (average pairwise similarity = 0.988). **(C)** Neighbor-Net phylogenetic network inferred in SplitsTree using p-distances. After excluding four long-branch isolates, the network resolved two primary clusters, designated Population 1 (blue branches) and Population 2 (red branches). Isolates are colored by geographic origin: United States (red), Australia (pink), United Kingdom (golden), Canada (blue), Poland (green), Norway (sky blue), and unknown origin (grey). Population 1 is predominantly composed of Australian isolates, whereas Population 2 includes isolates mainly from the United States, United Kingdom, Canada, and Norway. The Polish isolate SS62 clusters within Population 2 but is positioned near the boundary between the two populations.

Visual inspection of the phylogenetic tree revealed some host- and location-specific clustering, especially for clusters exhibiting slight enrichment for specific hosts (e.g., soybean) or locations (e.g., Ohio isolates). However, isolates from the same host or geographic origin were in general distributed throughout the tree rather than forming distinct monophyletic groups.

While *S. sclerotiorum* population structure models with K up to 5 were explored, only K = 2 and K = 3 produced consistent clustering patterns across replicates, whereas K = 4 and K = 5 failed to converge and did not yield reliable isolate assignments. The analysis supported K = 2 as the most biologically meaningful number of clusters based on the highest pairwise similarity across replicates. At K = 2, individuals were assigned to one of two distinct genetic groups, hereafter referred to as Population 1 and Population 2 (Fig. 2B). Population 1 was composed predominantly of isolates from Australia, with a small number of isolates from the US. In contrast, Population 2 included isolates primarily from the US, United Kingdom, Canada, and Norway. Replicate runs were highly consistent with an average pairwise similarity of 0.998, indicating stable clustering. When K was increased to 3, a third cluster appeared only at low ancestry proportions in a very small subset of isolates, and overall clustering consistency decreased with an average pairwise similarity of 0.988. This additional cluster did not correspond to a clearly differentiated population, suggesting over-partitioning rather than additional population structure.

Preliminary SplitsTree networks based on genome-wide SNV data identified four isolates with publicly available data (UM25, F20102, CU 11.14 and CU 11.21) that exhibited disproportionately long branch lengths, which obscured population-level structure. These isolates were excluded from subsequent network analyses to improve resolution. After filtering, the Neighbor-Net network clearly supported two genetic clusters, consistent with Population 1 and Population 2 previously identified (Fig. 2C). The network showed a high fit value (99.6%), indicating that the distances displayed in the split network closely reflected the original pairwise distances among isolates. Limited reticulation between the two populations further indicated strong genetic differentiation. This clustering pattern was consistent with the population structure previously inferred with fastSTRUCTURE, providing independent support for the presence of two major populations in the dataset. The Neighbor-Net network revealed a non-random geographic distribution, with Population 1 composed predominantly of Australian isolates, with a minor representation from the US and a single isolate from Norway, while Population 2 included isolates primarily from the USs, United Kingdom, Canada, and Norway. One isolate from Poland (SS62) clustered within Population 2 but was positioned near the boundary between the two populations, suggesting intermediate genetic placement.

After clone correction, the 216 isolates collapsed to 50 multilocus lineages in Population 1 (from 79 isolates) and 70 in Population 2 (from 137 isolates). Within population diversity was higher in Population 1 (π₁ = 0.022) than in Population 2 (π₂ = 0.017). The two populations were divergent, as the mean between population distance was d_xy_ = 0.028, and Nei’s net divergence was D_a_ = 0.0089 (95% bootstrap CI 0.0083–0.0096), which excludes zero and indicates divergence beyond shared ancestral variation. The distance-based F_ST_ was 0.31. AMOVA partitioned 50.8% of the total genetic variance among the two populations and 49.2% within them, yielding Φ_ST_ = 0.508 (*P* < 0.0001, 9,999 permutations).

### Phenotypic diversity

Among 114 US *S. sclerotiorum* isolates, substantial diversity was observed in radial growth measured as stAUGC, number of sclerotia produced per plate, sclerotial diameter, sclerotial weight, and number of days to initiation of sclerotia formation and sclerotia maturation (Supplementary Fig. 2). Values of stAUGC ranged from 11.5 to 63.6, with a median of 46.9 and interquartile range (IQR; Q1–Q3) of 38.1–57.5. The number of sclerotia produced per plate varied from 0 to 83, with a median of 46.5 and an IQR spanning 39 (Q1) to 58 (Q3). Sclerotial diameter varied from 1.57 to 3.47 mm, with a median of 2.48 and an interquartile range (Q1–Q3) of 2.27–2.78 mm. Similarly, sclerotial weight ranged from 0.95 to 8.72 mg, with a median of 3.81 and an interquartile range of 2.74–5.15 mg. The time to sclerotial initiation ranged from 3 to 10 days, with a median of 4 days and an interquartile range (Q1–Q3) of 3–4 days. Days to sclerotial maturation ranged from 4 to 11 days, with a median of 6 and an interquartile range of 5–6. Isolates SS166 and SS167 did not grow on PDA and therefore did not produce sclerotia. SS158 initiated sclerotia formation but failed to successfully develop them. Consequently, no sclerotial data other than days to formation of sclerotia were collected for this isolate, and values were recorded as zero in the box plots.

Principal component analysis after removing outlier isolates that did not grow well on PDA (SS158, SS166, and SS167) revealed that the first two principal components collectively explained 66.9% of the total variance (PC1: 39.1% and PC2: 27.8%; Fig 3A and 3B). PC1 represented a trade-off between radial growth rate and sclerotial development timing (sclerotial formation and maturation), where it separated isolates exhibiting rapid radial growth with shorter sclerotial development time from those with slow growth but longer sclerotial formation time. Highest PC1 isolates (SS161, SS213, SS165, and SS23) invested substantially in sclerotial formation with correspondingly slow radial growth. In contrast, the lowest PC1 isolates (SS121, SS137, and SS113) prioritized rapid radial growth with shorter sclerotial development time. PC2, on the other hand, captured trade-off between sclerotial number and weight and diameter, contrasting isolates producing numerous small sclerotia with those producing fewer larger sclerotia. Along PC2, isolates with high values (SS214, SS187, and SS113) produced numerous small sclerotia, whereas low PC2 isolates (SS218, SS43, SS195, and SS169) produced fewer but larger sclerotia. However, isolates within each PC group did not necessarily share a common host or geographical location.

**Figure 3.**
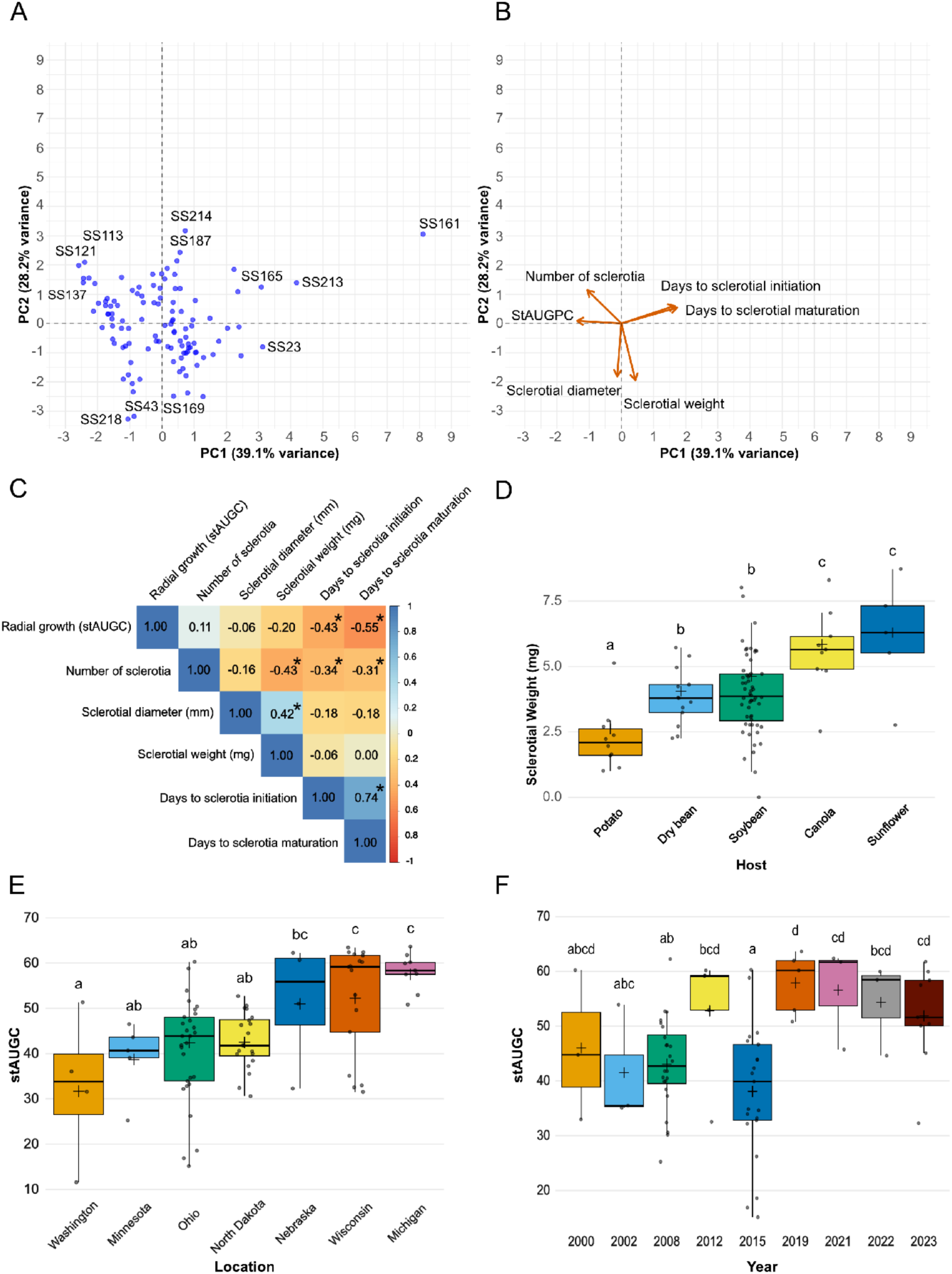
**(A)** Principal component analysis (PCA) of *Sclerotinia sclerotiorum* isolates after excluding poorly performing outlier isolates (SS158, SS166, and SS167) on potato dextrose agar (PDA). The first two principal components explained 66.9% of the total variance (PC1: 39.1%; PC2: 27.8%). The plot shows the distribution of isolates, with each point representing an individual isolate. **(B)** Loading plot showing the contribution of six morphological traits to PC1 and PC2. **(C)** Correlation matrix illustrating the relationships among the six morphological traits. Values represent Pearson’s correlation coefficients, with color intensity indicating the strength and direction of the associations. An asterisk (*) denotes statistical significance at *P* < 0.01 **(D)** Host significantly affected sclerotial weight (*P* = 2.4 × 10⁻^5^). Sunflower and canola produced the heaviest sclerotia and did not differ from each other, followed by soybean and dry bean, while potato isolates produced the lightest sclerotia. Data are presented as box plots, where the central line represents the median, the “+” symbol indicates the mean, and boxes and whiskers show the distribution of the data. **(E)** Geographic location significantly affected radial colony growth, measured as the standardized area under the growth curve (stAUGC) (*P* = 1.0 × 10⁻^4^). Estimated marginal means showed the highest values for Michigan, followed by Wisconsin and Nebraska, while Washington had the lowest and Minnesota, Ohio, and North Dakota exhibited intermediate values. Data are presented as box plots, where the central line represents the median, the “+” symbol indicates the mean, and boxes and whiskers show the distribution of the data. **(F)** Year of collection significantly affected radial colony growth, measured as the standardized area under the growth curve (stAUGC) (*P* = 0.01). Values were generally higher in recent years (2019-2023) and lowest in 2015, though most pairwise comparisons were not significant. Data are presented as box plots, where the central line represents the median, the “+” symbol indicates the mean, and boxes and whiskers show the distribution of the data.

### Effect of host, location, and year on phenotypic traits

To formally test if host, location, and year had effects on phenotypic traits, phylogenetic linear mixed-effects models were fitted to take into account isolate relatedness. Prior to modeling, Pearson’s correlation analysis revealed several significant associations among morphological and cultural traits of *S. sclerotiorum*, as the PCA analyses suggested (Fig. 3C). Median stAUGC showed a significant negative correlation with days to sclerotial initiation (r = −0.43, *P* = 2.4 × 10⁻⁶) and days to sclerotial maturation (r = −0.55, *P* = 5 × 10⁻^10^). The number of sclerotia per plate was significantly negatively correlated with sclerotial weight (r = −0.43, *P* = 2.6 × 10⁻⁶), days to sclerotial initiation (r = −0.34, *P* = 2.1 × 10⁻^4^), and days to sclerotial maturation (r = −0.31, *P* = 8.8 × 10⁻^4^). Sclerotial diameter showed a significant positive correlation with sclerotial weight (r = 0.42, *P* = 5.1 × 10⁻⁶). Days to sclerotial initiation and days to sclerotial maturation were positively correlated (r = 0.74, *P* = 2.2 × 10⁻^1^⁶). Because several traits were significantly correlated, we selected stAUGC and sclerotial weight as representative variables for subsequent modeling. The remaining co-varying traits were excluded to reduce collinearity.

The host had no effect on stAUGC (*P* = 0.76). Radial growth production of the isolate did not differ among host categories. In contrast, host had a significant effect on sclerotial weight (*P* = 2.4 × 10⁻^5^). Mean sclerotial weight differed among host groups (pairwise comparisons, α = 0.05), isolates from sunflower (6.29 mg) and canola (5.85 mg) did not differ from each other but produced heavier sclerotia than isolates from soybean (4.62 mg) and dry bean (4.06 mg), while isolates from potato (2.61 mg) produced significantly lighter sclerotia than all other host groups (Fig. 3D; Supplementary Table 1).

Geographic location had a significant effect on radial growth (stAUGC; *P* = 1.0 × 10⁻^4^). Estimated marginal means indicated substantial variation among locations, with isolates from Michigan exhibiting the highest stAUGC values (57.5), followed by Wisconsin (52.2) and Nebraska (51.0) (Fig. 3E; Supplementary Table 2). In contrast, isolates from Washington showed the lowest stAUGC (31.7), while Minnesota (38.7), Ohio (42.4), and North Dakota (42.5) exhibited intermediate values. No significant effect of geographic location was detected for sclerotial weight (*P* = 0.38), indicating that sclerotial weight did not vary consistently among locations.

Year of collection had a significant effect on stAUGC (*P* = 0.01). Estimated marginal means indicated substantial interannual variation in stAUGC, with higher values observed in more recent years (2019-2023) and lower values in earlier years, particularly 2015. However, pairwise comparisons revealed extensive overlap among years, with most year-to-year contrasts not statistically significant. Based on pairwise comparisons, 2015 formed the lowest grouping with a mean of 38.1, while 2019 formed the highest grouping (57.9) and the remaining years exhibited intermediate, overlapping groupings (Fig. 3F; Supplementary Table 3). On the other hand, no significant effect of year of collection was detected for sclerotial weight (*P* = 0.49), indicating that sclerotial weight did not vary consistently across years.

### Phylogenetic signal analyses

To explore potential associations between genetic variation, phenotypic variation, and associated metadata among *S. sclerotiorum* isolates, we applied Blomberg’s K and Pagel’s λ to assess phylogenetic signals of measured phenotypes and collection metadata. Blomberg’s K and Pagel’s λ revealed generally weak associations between the phylogeny and the measured phenotypic traits (Table 2). However, relatively higher Pagel’s λ values were observed for sclerotial weight and sclerotial initiation compared to other phenotypic traits. In contrast, phylogenetic diversity appears to show strong association with host, geographic origin, and year of collection based on higher Pagel’s λ values estimated as a tree-transformation parameter via likelihood optimization of Mk model for categorical traits indicating that these factors are associated with the genetic structuring of populations (Table 2). The phylogeny of the isolates, aligned with a heatmap of phenotypic traits and isolate metadata (host, geographic origin, and year of collection), is shown in Fig. 4. Closely related isolates tended to originate from the same host, location, and collection period, whereas phenotypic traits, including radial colony growth (stAUGC), number of sclerotia per plate, sclerotial diameter, sclerotial weight, and days to sclerotial initiation and maturation, were distributed across the phylogeny, with high and low trait values occurring throughout rather than concentrated within particular clades.

**Figure 4.**
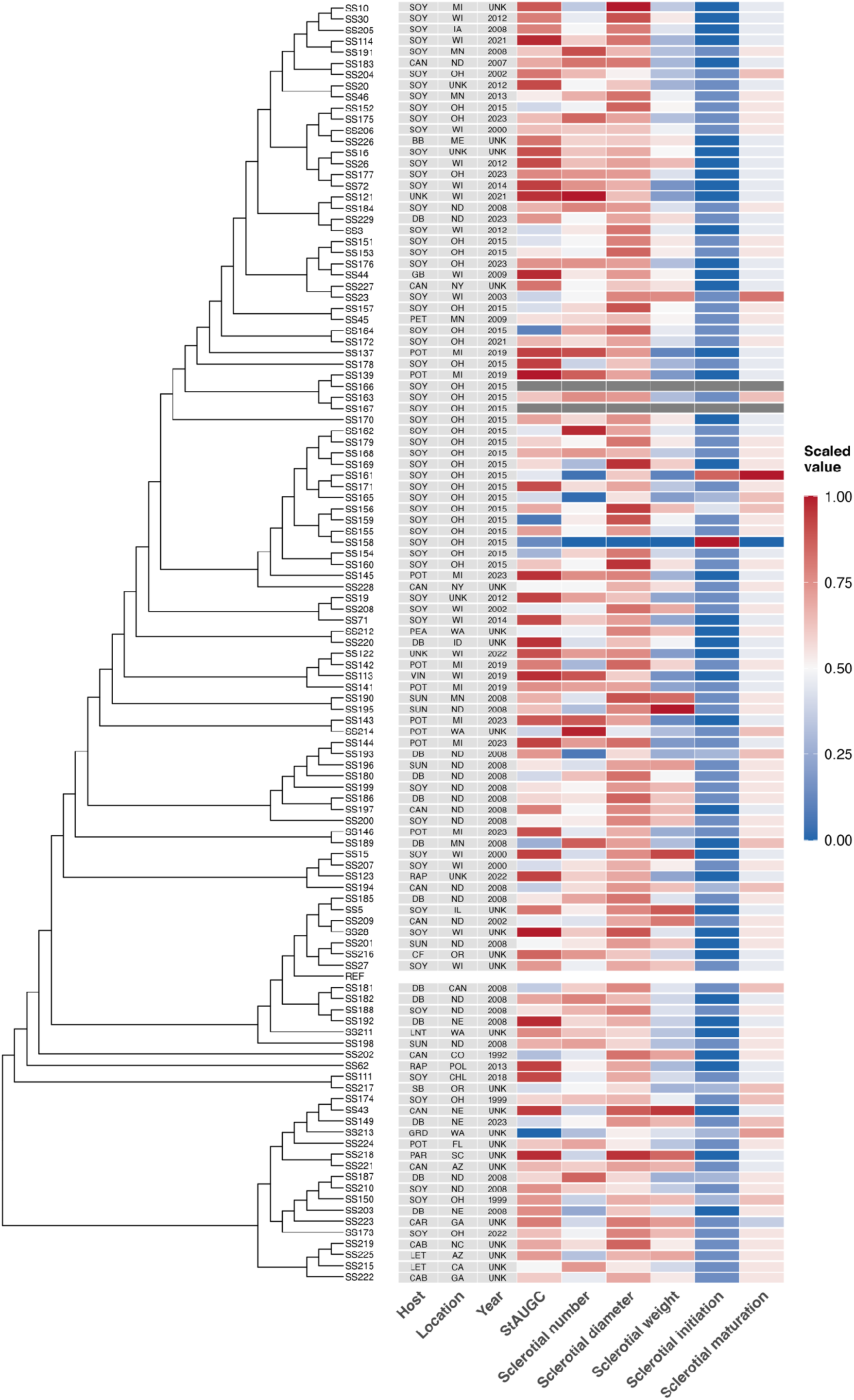
Phylogenetic associations among *Sclerotinia sclerotiorum* isolates shown alongside a heatmap of phenotypic traits and associated metadata, including geographic origin, host, and year of collection. The combined visualization illustrates potential associations between genetic variation, morphological traits, and isolate metadata. Color intensity represents relative trait values, with red indicating higher values and blue indicating lower values. Host abbreviations: BB, bush bean; CAB, cabbage; CAN, canola; CAR, carrot; CF, cauliflower; DB, dry bean; GB, green bean; GRD, gourd; LET, lettuce; LNT, lentil; PAR, parsley; PEA, pea; PET, petunia; POT, potato; RAP, rapeseed; SB, snap bean; SOY, soybean; SUN, sunflower; VIN, vinca (periwinkle). Locations are given as two-letter US state postal codes, except CAN, Canada; CHL, Chile; and POL, Poland. UNK, unknown host, location, or year. stAUGC, standardized area under the growth curve for radial colony growth.

**Table 2.** Phylogenetic signal estimates using Blomberg’s K and Pagel’s λ for numerical traits and Pagel’s λ estimated as a tree-transformation parameter via likelihood optimization of Mk model for categorical traits.

| **Response variable** | **Blomberg’s K** | **Pagel’s λ** |
|---|---|---|
| stAUGC<sup>a</sup> | 0.079 | 0.00007 |
| Sclerotia number | 0.084 | 0.066 |
| Sclerotia diameter | 0.077 | 0.00007 |
| Sclerotia weight | 0.097 | 0.483 |
| Sclerotial initiation | 0.090 | 0.231 |
| Sclerotial maturation | 0.074 | 0.00007 |
| Host | - | 0.893<sup>*</sup> |
| Location | - | 0.899<sup>*</sup> |
| Year | - | 0.744<sup>*</sup> |

<sup>a</sup> Radial colony growth measured as the standardized area under the growth curve (stAUGC).
<sup>\*</sup> Strong phylogenetic signal indicating that closely related isolates tend to share the same host, geographic origin and year of collection.

## Discussion

Broad host range pathogens present a particular challenge for understanding population structure, because their ability to infect many hosts across diverse environments may obscure associations between genetic relatedness, host, and geographic origin. *S. sclerotiorum* is an ideal system for examining these relationships. The remarkable success of *S. sclerotiorum* as a pathogen is reflected in its ability to infect over 400 plant species worldwide, making it one of the most versatile fungal pathogens (Boland and Hall 1994). However, its population structure remains incompletely understood, in part because of its mixed reproductive strategy, combining homothallic sexual reproduction with extensive clonal spread, complicates interpretation of genetic exchange within and among populations (Attanayake et al. 2019; Buchwaldt et al. 2022). Our study is among the few studies to examine the genetic diversity of *S. sclerotiorum* across a wide range of host crops, a broad geographic area, and multiple years of isolation in the US. Here, we characterized genetic variation and inferred evolutionary relationships through phylogenetic and population genetic analysis. This approach revealed substantial genetic diversity ranging up to 0.06 substitutions per site among isolates (Fig. 2A). This finding may suggest that genetic variation accumulated within the population over time through mutation and recombination, resulting in multiple genetically differentiated subpopulations rather than a homogeneous population, potentially driven by adaptation to distinct ecological niches or geographic regions. Despite this overall diversity, the US population is characterized by a striking preponderance of tightly clustered isolates with short branch lengths (< 0.01 substitutions per site), suggesting potential clonal relationships within these groups. This dual pattern of substantial overall divergence coupled with clonal clustering indicates a complex evolutionary history shaped by ancient divergence events while maintaining local population structure, a finding that enhances our understanding of *S. sclerotiorum* population genetics in North America. Consistent with other fungi, such as *Puccinia striiformis* f. sp. *tritici* and *Cryphonectria parasitica*, that reproduce sexually in their native range but spread as clonal lineages, reflecting the view that successful clones emerge from recombining populations (Ali et al. 2014; Drenth et al. 2019; Milgroom et al. 2008).

Population structure and phylogenetic network analyses consistently supported the presence of two major genetic populations at the global scale (Fig. 2B and 2C). Population 1 consisted predominantly of Australian isolates, whereas Population 2 contained most isolates from North America and Europe, including the US, United Kingdom, Canada, and Norway. The composition of these populations may reflect adaptation to distinct climatic conditions following introduction into different geographical regions. Australia is characterized largely by warmer and drier environments with seasonal drought compared to the cooler temperate climates with seasonal interruption due to cold or freezing winters represented by many North American and European locations. Over time, local environmental selection pressures may have contributed to the divergence observed between the two populations. Similar observations have been reported previously that adaptation to distinct environmental conditions may have shaped divergence between the populations (Derbyshire et al. 2019a). Nevertheless, neither population was entirely geographically exclusive. The presence of several US isolates within Population 1, which is predominantly Australian cluster, suggests inter-continental long-distance dispersal or historical movement of infected plant material, which may also facilitate the admixture between populations. Such movement is plausible given the broad host range of *S. sclerotiorum* and the international trade of agricultural commodities. Evidence of admixture between European-North American and Australian-African populations has also been documented recently (Derbyshire et al. 2025). Moreover, the US isolates associated with the Australian-dominated cluster were not closely related to one another within the network, but instead were distributed across different branches of Population 1 and originated from different regions of the US. This pattern suggests that these isolates likely represent multiple independent introduction events rather than a single transmission event followed by local spread.

Consistent with multiple independent introductions into Population 1, the greater within population nucleotide diversity of Population 1 relative to Population 2 (π₁ > π₂) may reflect the introduction and persistence of genetically distinct lineages arising from such dispersal events. Mean between population divergence (D_xy_ = 0.028) and Nei’s net divergence (D_a_ = 0.0089; 95% CI: 0.0083–0.0096) indicate substantial differentiation beyond shared ancestral variation, suggesting that the populations have experienced evolutionary separation while still retaining signatures of historical exchange. The distance-based F_ST_ value (0.31) further supports moderate-to-strong differentiation between populations. Consistent with these patterns, AMOVA demonstrated that 50.8% of the total genetic variation was partitioned among populations, while 49.2% occurred within populations, resulting in a significant Φ_ST_ value (0.508, *P* < 0.0001; 9,999 permutations). These results confirm strong population structure while also indicating substantial within-population variation. Taken together, these findings suggest that the observed population structure has been shaped by historical intercontinental movement facilitating admixture between geographically distinct regions, in combination with long-term population differentiation.

*S. sclerotiorum* isolates examined in this study exhibited remarkable phenotypic diversity across all morphological and developmental traits. The first principal component captured a trade-off between radial growth and sclerotial development timing (39.1%), and the second principal component revealed a further trade-off between sclerotial number and sclerotial size (27.8%) (Fig. 3A and 3B). This observation indicates that the diversity of certain phenotypic features is structured rather than random and raises important questions about the ecological and evolutionary pressures shaping these traits and their potential relevance to disease management. The radial growth strategy may reflect adaptation to host environments where rapid colonization of tissue is advantageous, such as in senescing hosts, whereas the sclerotial development strategy may be more beneficial in environments where persistence and survival between host crops is paramount. The trade-off between sclerotial number and sclerotial size could be advantageous under different circumstances and ecological contexts (Sharma et al. 2026a). Numerous small sclerotia may maximize propagule density and dispersal potential while being less likely to be removed with the grain, remaining in the field to contribute to future inoculum loads (Webster et al. 2025). This strategy may be particularly advantageous in cropping systems where sclerotia can germinate directly into mycelia and establish infection. Additionally, biological control agents (such as *Coniothyrium minitans*) that target the sclerotial stage may be less effective against isolates producing numerous small sclerotia, which are more cryptic in soil and therefore more difficult for biocontrol organisms to locate and colonize compared to larger, more conspicuous sclerotia (Sharma et al. 2026b). Conversely, fewer larger sclerotia may represent an allocation strategy favoring quality over quantity, with the assumption that larger sclerotia possess more substantial reservoir for apothecial germination and apothecia production (Sharma et al. 2026a; Taylor et al. 2018). This is particularly important in crops where ascospores are the major source of infection, as larger sclerotia with greater energy reserves would support the development and production of apothecia and the subsequent release of infectious ascospores. The phenotypic groups identified through PCA did not seem to correspond to a single geographic origin or host source, suggesting that phenotypic diversity is maintained within populations rather than being geographically or host partitioned. This observation raises the possibility that diverse phenotypes confer fitness advantages under varying environmental conditions within a single geographic or host context, or alternatively, that microhabitat variation has resulted in the maintenance of multiple phenotypes without contemporary selection for differentiation.

We further investigated whether host, location, and year of collection affected individual phenotypic traits using phylogenetic linear mixed-effects models. The host plant was significantly associated with sclerotial weight, with isolates from sunflower and canola producing the heaviest sclerotia, whereas isolates from soybean and dry bean produced significantly lighter sclerotia (Fig. 3D). Most strikingly, potato isolates produced substantially lighter sclerotia than all other host groups. This could be because different hosts may vary in nutritional quality or total biomass availability. Similar trends have been reported previously, where isolates produced heavier sclerotia on oilseed rape than on potato (Taylor et al. 2018). Geographic location was significant associated with radial growth (stAUGC), with isolates from Michigan exhibiting the highest rates, followed by Wisconsin and Nebraska, while isolates from Washington displayed markedly lower rates (Fig. 3E). This could be due to regional differences in growing conditions, including temperature, humidity, and moisture patterns, which may select for locally adapted phenotypes, with cooler or more favorable growing regions (such as the northern Great Lakes) supporting the persistence of faster-growing isolates. Alternatively, regional differences in agricultural practices, crop rotations, or cropping history may influence pathogen phenotypic composition. The reduction in radial growth in Washington isolates warrants investigation; this could reflect geographic isolation of Washington populations or environmental selection pressures specific to Pacific Northwest growing conditions. Population differences associated with hot and cool climates in the US have also been reported previously in population genetic studies (Poudel et al. 2023). Year of collection was significant associated with radial growth (stAUGC), with a general trend toward higher growth rates in more recent years (2019-2023) compared to earlier years, particularly 2015 (Fig. 3F). Annual variation in weather conditions, host type, and micro-environment during that or previous growing season could influence pathogen phenotype expression. It is also possible that the maintenance of isolates in the laboratory has influenced phenotypic expression, with differences in storage time or re-culturing history between early and recent isolates affecting radial growth rates independent of original phenotype.

Our analyses employed Blomberg’s K and Pagel’s λ to quantify phylogenetic signal across phenotypic traits and associated metadata, revealing patterns that fundamentally distinguish between traits influenced by evolutionary history versus those shaped by contemporary ecological and demographic factors. We observed a weak phylogenetic signal across most phenotypic traits measured in this study, suggesting that phenotypic variation in these traits is not primarily structured by the evolutionary relationships among isolates. This may indicate that these traits are environmentally plastic, with variation influenced more by microhabitat conditions and environmental stress factors than by shared ancestry. Alternatively, if these traits were controlled by only a few genes with large effects, we would expect closely related isolates to show more similar phenotypes and therefore exhibit a stronger phylogenetic signal. Instead, the weak phylogenetic signal suggests that phenotypic variation is likely governed by many genes with small effects, consistent with proteomic evidence showing that sclerotial development involves numerous proteins spanning multiple functional categories (Liang et al. 2010). In contrast, associated metadata representing host species, geographic origin, and year of collection showed a strong phylogenetic signal, indicating that phylogenetic diversity is strongly associated with these ecological and demographic variables. Although *S. sclerotiorum* has a relatively closed pan-genome and host specialization appears unlikely (Derbyshire et al. 2025), the up-regulation of small RNAs involved in silencing host mRNAs through complementary base pairing has been reported to include both conserved and host-specific responses (Derbyshire et al. 2019b). This may reflect enhanced compatibility with particular hosts driven by coevolutionary dynamics, in which pathogen lineages track host availability or host resistance evolution, ultimately contributing to host-associated genetic structure. Geographic clustering in the phylogeny can arise from independent introductions of the pathogen into different regions, producing founder effects. Subsequent regional diversification and limited gene flow caused by spatial separation can lead to the accumulation of distinct mutations over time, ultimately resulting in phylogenetic clustering by geographic origin. For example, populations of *S. sclerotiorum* from geographically distant Canadian provinces showed greater genetic differentiation and lower gene flow than populations from neighboring provinces, indicating that spatial separation restricted gene exchange (Buchwaldt et al. 2022). We speculate that the tendency for isolates collected in the same year to be more closely related to one another than to isolates from other years may reflect selection favoring advantageous traits such as more aggressive or persistent isolates in field populations, driven by weather conditions, agricultural practices, and fungicide use (Derbyshire 2020).

Taken together, our study has important implications for understanding the evolution and adaptation of *S. sclerotiorum*, which in turn informs our knowledge of disease epidemiology and the development of effective management strategies. The evidence of historical intercontinental movement between US and Australian populations highlights the importance of understanding global pathogen dispersal patterns for biosecurity and quarantine decisions. The discovery of considerable genetic and phenotypic diversity among US populations suggests that disease management strategies may need to be tailored differently across regions. Moreover, the observed associations between phylogeny and host, geographic origin, and collection time indicate that genomic characterization can provide valuable information about pathogen population dynamics beyond observable phenotypic traits alone. Collectively, these findings establish a foundation for future surveillance efforts and may help inform resistance breeding programs aimed at developing crop cultivars with improved durability against diverse pathogen populations.

## Data Availability Statement

Newly generated sequencing reads have been deposited under BioProject PRJNA1552207. Publicly available sequence data were obtained from NCBI under BioProject PRJNA1120954 (Derbyshire et al. 2025).

## Acknowledgments

We gratefully acknowledge the Wisconsin Soybean Marketing Board (WSMB) and the North Central Soybean Research Program (NCSRP) for providing partial support for this research. We thank Drs. Anne Dorrance and Mitchell Roth for preserving and recovering historical *Sclerotinia sclerotiorum* isolates and the current Soybean Pathology and Nematology team at The Ohio State University for collecting and maintaining the isolate collection. H.D.L.N. received support from the Ohio Soybean Council (Project 23-R-29) and the USDA National Institute of Food and Agriculture (Hatch Project OHO01569). E.G.-T and S.O. are supported by the Office of the Vice Chancellor for Research and Graduate Education at the University of Wisconsin-Madison with funding from the Wisconsin Alumni Research Foundation and the Department of Plant Pathology at the University of Wisconsin-Madison.

## CRediT Author Contributions

A.S.: Conceptualization, Methodology, Data curation, Formal analysis, Writing – original draft; C.L.G.: Methodology; Writing – review & editing; S.O.D.: Methodology, Writing – review & editing; R.W.W.: Resources, Writing – review & editing; H.D.L.N.: Resources, Writing – review & editing; D.L.S.: Conceptualization, Methodology, Writing – review & editing, Project administration, Supervision; E.G.T.: Conceptualization, Methodology, Writing – review & editing, Project administration, Supervision.

---

**Supplementary Figure. 1.**
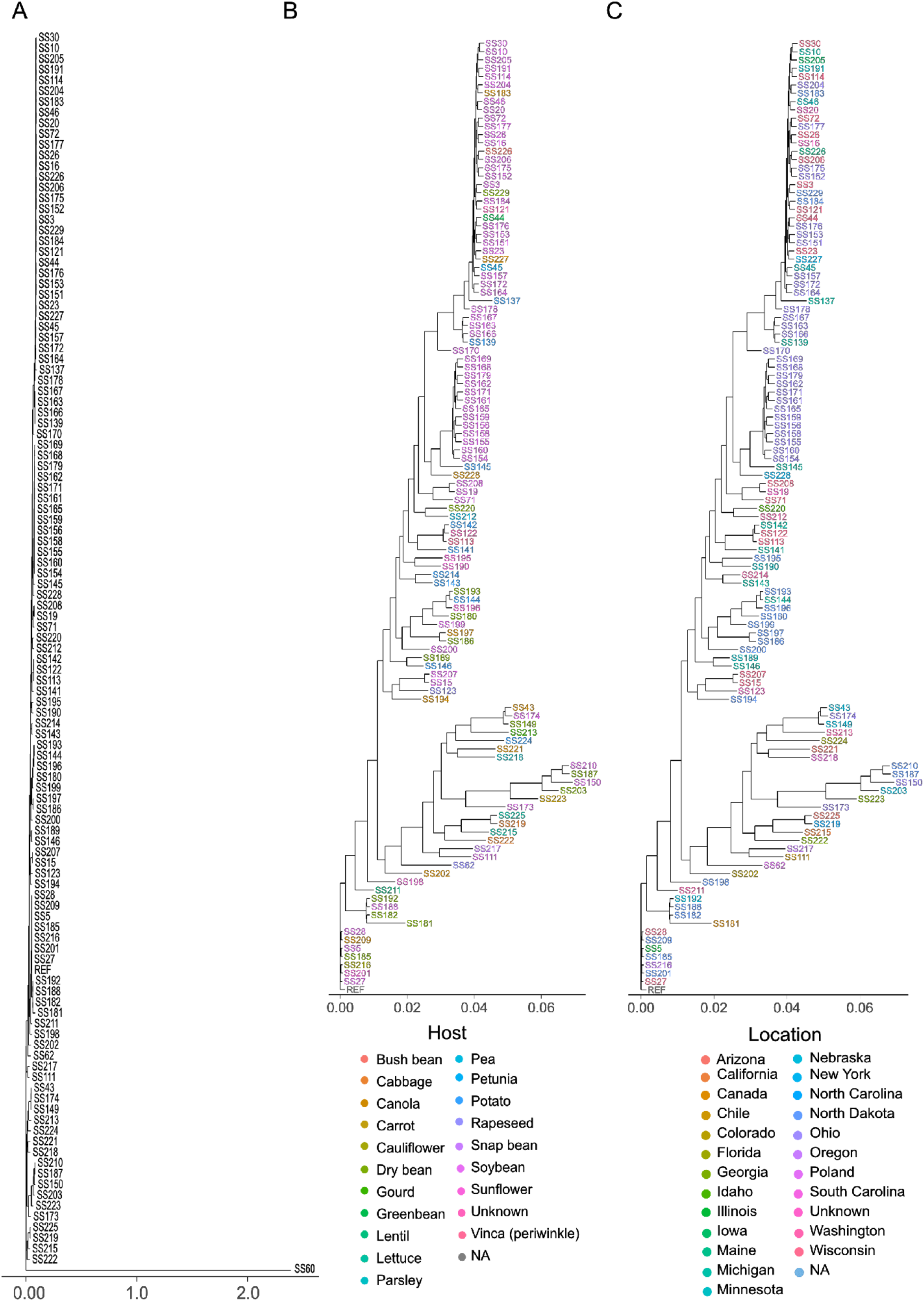
**(A)** Maximum-likelihood phylogenetic tree of newly sequenced *Sclerotinia sclerotiorum* isolates based on whole genome-wide SNP analysis. The tree was rooted with the outgroup isolate SS60 to provide evolutionary context and assess divergence among isolates. Branch lengths are proportional to genetic distance. **(B)** Maximum likelihood phylogenetic tree with isolates colored according to host plant origin. The tree shows limited host-specific clustering; however, some clusters exhibit slight enrichment for particular hosts (e.g., soybean). **(C)** Maximum likelihood phylogenetic tree with isolates colored by geographic origin. The tree shows limited location-specific clustering; however, some clusters exhibit slight enrichment for particular locations (e.g., isolates from Ohio).

**Supplementary Figure. 2.**
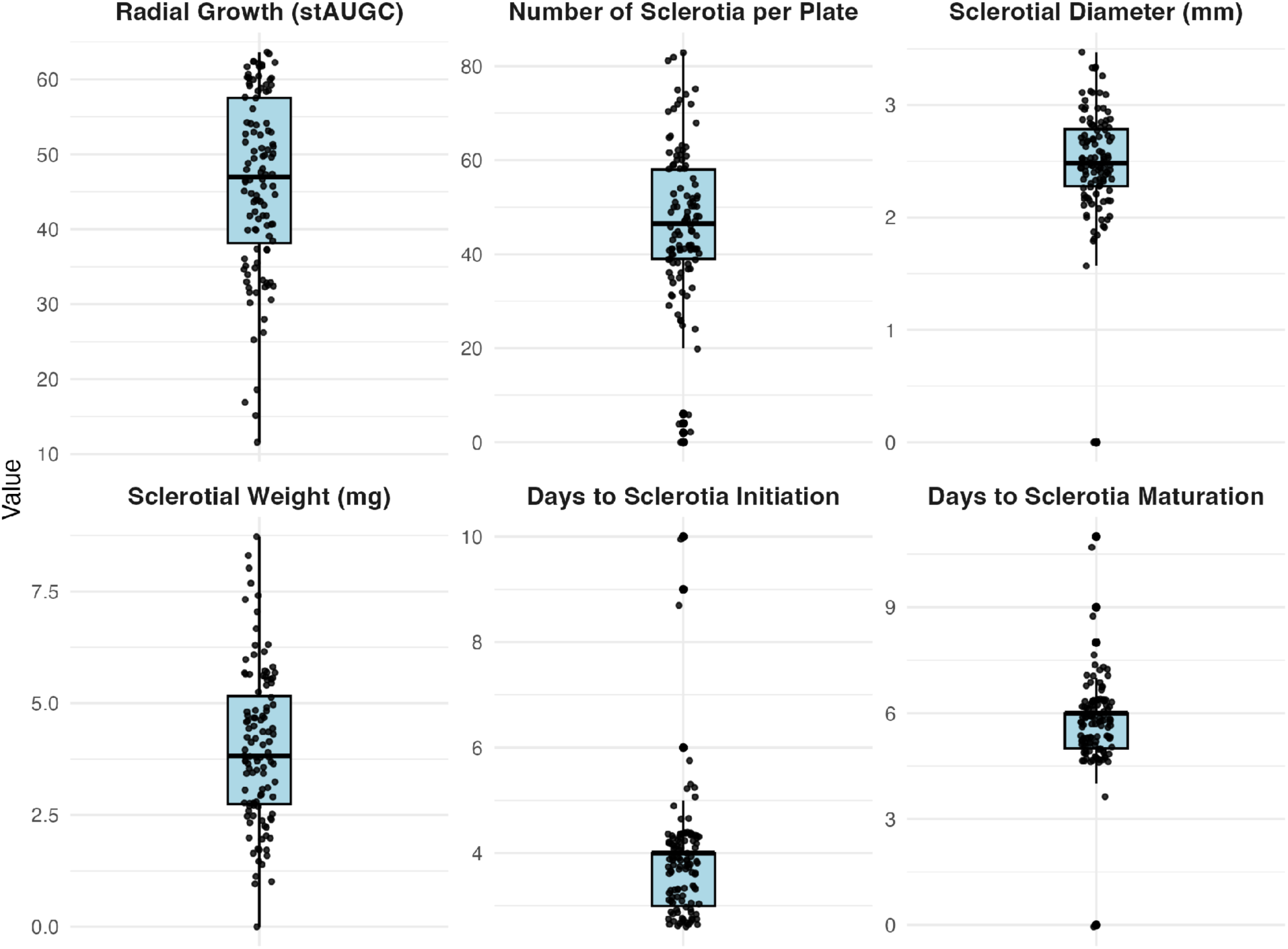
Morphological characteristics of *Sclerotinia sclerotiorum* isolates grown on PDA. Box plots represent variation in radial colony growth, measured as the standardized area under the growth curve (stAUGC), number of sclerotia per plate, sclerotial diameter (mm), sclerotial weight (mg), days to sclerotia initiation, and days to sclerotia maturation across isolates. Boxes indicate interquartile ranges, horizontal lines denote medians, and whiskers represent minimum and maximum values.

**Supplementary Table 1.** Estimated marginal means of median sclerotial weight for *Sclerotinia sclerotiorum* isolates by host crop. Different letters indicate significant differences among hosts based on pairwise comparisons of estimated marginal means (*P* < 0.05).

| **Host** | **Mean (sclerotial weight)** | **Standard error** | **Group** |
|---|---|---|---|
| Potato | 2.61 | 0.61 | a |
| Dry bean | 4.06 | 0.56 | b |
| Soybean | 4.62 | 0.46 | b |
| Canola | 5.85 | 0.59 | c |
| Sunflower | 6.29 | 0.78 | c |

**Supplementary Table 2.** Estimated marginal means of median standardized area under the growth curve (stAUGC) for *Sclerotinia sclerotiorum* isolates by geographical location. Different letters indicate significant differences among locations based on pairwise comparisons of estimated marginal means (*P* < 0.05).

| **Location** | **Mean (stAUGC)** | **Standard error** | **Group** |
|---|---|---|---|
| Washington | 31.7 | 5.54 | a |
| Minnesota | 38.7 | 4.76 | ab |
| Ohio | 42.4 | 2.50 | ab |
| North Dakota | 42.5 | 2.96 | ab |
| Nebraska | 51.0 | 5.23 | bc |
| Wisconsin | 52.2 | 2.59 | c |
| Michigan | 57.5 | 3.68 | c |

**Supplementary Table 3.**
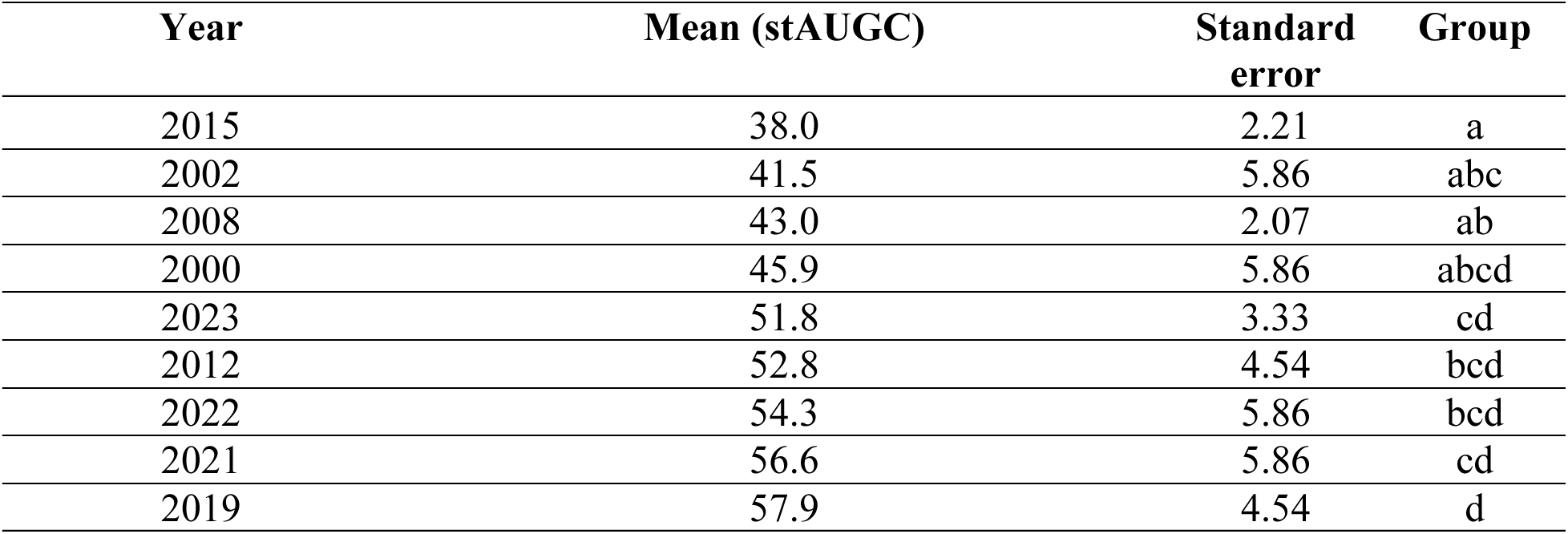
Estimated marginal means of median standardized area under the growth curve (stAUGC) for *Sclerotinia sclerotiorum* isolates by year of collection. Different letters indicate significant differences among years based on pairwise comparisons of estimated marginal means (*P* < 0.05).

| **Year** | **Mean (stAUGC)** | **Standard error** | **Group** |
|---|---|---|---|
| 2015 | 38.0 | 2.21 | a |
| 2002 | 41.5 | 5.86 | abc |
| 2008 | 43.0 | 2.07 | ab |
| 2000 | 45.9 | 5.86 | abcd |
| 2023 | 51.8 | 3.33 | cd |
| 2012 | 52.8 | 4.54 | bcd |
| 2022 | 54.3 | 5.86 | bcd |
| 2021 | 56.6 | 5.86 | cd |
| 2019 | 57.9 | 4.54 | d |

